# Feasibility study of fluid-structure-growth modeling in the context of endogenous tissue restoration

**DOI:** 10.64898/2026.10.20.760334

**Authors:** Lauranne Maes, Amber Hendrickx, Hadi Shirazi, Deniz Ozturk, Maria Bastron, Tahir Turgut, Vincent Bouwman, Nils Götzen, Thibault Vervenne, Malte Rolf, Felipe Sempértegui, Martijn Cox, Stéphane Avril, Gerhard A. Holzapfel, Bart Meuris, Nele Famaey

## Abstract

*In situ* tissue engineering exploits natural inflammatory responses to turn a synthetic vascular graft into a regenerated, functioning, and living vessel through endogenous tissue restoration. Despite promising preclinical results, challenges remain, especially in high-pressure regimes. To better understand the underlying biological processes and predict long-term outcomes of graft implantation, we propose an *in silico* approach. We account for the solid mechanics of conduit grafts in high and low pressure regimes, the wall shear stresses, and long-term growth and remodeling through two reduced-order fluid-structure-growth models. The first employs an artificial neural network surrogate model to compute the time-averaged wall shear stress at every growth step, whereas the second evaluates a fully-coupled fluid-structure interaction model every 60 time steps. We thoroughly verify both models and confirm acceptable model-form errors, then calibrate them against data from animal experiments. A validation study comparing *in silico* predictions with unseen animal data of carotid and pulmonary conduit grafts confirms that the model remains reliable up to at least one month of prediction time, hence in the future enabling *in silico* optimization of short-term endogenous tissue restoration in tissue engineering vascular grafts.

## 1 Introduction

*In situ* tissue engineering exploits the body’s natural inflammatory response so that a biodegradable synthetic cardiovascular implant gradually transform into a regenerated, functioning, and living blood vessel or heart valve, a process called endogenous tissue restoration (ETR). The technique has shown successful outcomes in various preclinical large animal models (Fukunishi et al., 2017; Drews et al., 2020) as well as in clinical trials on pediatric patients with congenital malformations, specifically for pulmonary valve replacement (Morales et al., 2021) and venous bypassing (Hibino et al., 2010; Bockeria et al., 2020). However, these results stem from low-pressure applications, and challenges remain in high-pressure regimes, where aneurysm formation alongside adverse tissue remodeling has been reported (Duijvelshoff et al., 2021).

ETR is generally described as an inflammatory phase with immune cell infiltration, followed by a proliferative phase with secondary cell recruitment and re-endothelialization, and finally an extracellular matrix (ECM) remodeling phase leading to tissue homeostasis (Wissing et al., 2017). However, the complex interplay of cellular infiltration, recruitment, ECM production, and remodeling remains poorly understood, and optimal implant designs are therefore still lacking.

Thus, we propose an *in silico* approach to improve the understanding of these processes and to predict outcomes of graft implantation for different geometries, graft locations, and boundary conditions.

Previously, Wissing et al. (2020) showed that wall shear stress likely plays a key role in ETR. Many computational models of ETR, including our previous work (Shirazi et al., 2025), simplify its calculation by assuming idealized Hagen-Poiseuille flow (Szafron et al., 2018; Drews et al., 2020), while Latorre et al. (2022) proposes a reduced-order one-dimensional flow model. Fluid-structure-growth (FSG) models resolving the three-dimensional Navier-Stokes equations have been developed mainly in the context of aneurysm growth (Figueroa et al., 2009; Watton et al., 2009; Aparício et al., 2014; Grytsan et al., 2015; Teixeira et al., 2020; Mousavi et al., 2021), although Schwarz et al. (2023) propose a fully coupled FSG solver for tissue-engineered vascular grafts. Comparing it against the Hagen-Poiseuille assumption, they demonstrate the importance of resolving the three-dimensional fluid field.

Such fully coupled FSG models remain computationally expensive. We therefore develop two reduced-order FSG approaches, both using the homogenised constrained mixture theory (Cyron et al., 2016) for the growth and remodeling part, combined with two different flow models. In the first, a low-fidelity flow model continuously informs the growth and remodeling model. In the second, a high-fidelity flow model informs the growth and remodeling model at sparser time intervals. We thereby compare two opposite reduced-order approaches and examine the feasibility of these models for their use in the context of *in situ* tissue engineering at reasonable computational cost. We present the models in Section 2.2 and thoroughly verify (Section 3.2), calibrate (Section 3.1), and validate (Section 3.3) them against the experimental data outlined in Section 2.1. Compared to our previous study (Shirazi et al., 2025), a more thorough validation is performed, now considering an extra set of animal experiments. Moreover, the model’s behaviour against experimental data is assessed in both high-pressure and low-pressure regimes, now considering both carotid and pulmonary conduit grafts.

## 2 Methods

### 2.1 Animal experiments

Experimental data for model validation were obtained for two graft types, a carotid conduit graft and a pulmonary valved conduit graft. The carotid graft experiments were approved by the Animal Ethics Committee of KU Leuven (no. P093/2020). Ten biodegradable carotid artery grafts of two designs were implanted in sheep. Six animals received a thicker graft with a wall thickness of 0.63 mm, and four received a thinner wall 0.40 mm. All grafts had an initial diameter of 6.0 mm and a length of 70.0 mm, and the follow-up period was six months.

Under a separate protocol approved by the same committee (no. P189/2021), the pulmonary valved conduit experiments were carried out, in which biodegradable pulmonary valved conduits of two designs were implanted in eight sheep. Four animals received valves with a wall thickness of 0.70 mm and the remaining four animals received valves with a thinner wall of 0.57 mm. All valve grafts had a diameter of 20.0 mm, while the length was anatomy-dependent and ranged from 22.7 to 31.8 mm. Only the conduit segment distal to the valve was considered for the validation of the computational models described below.

Carotid and pulmonary graft diameters were evaluated by ultrasound imaging at multiple time points throughout the follow-up period, at approximately 15, 30, 60, 90, 120, 150, and 180 days for the carotid grafts, and approximately 21, 60, 120, and 180 days for the pulmonary grafts. Flow measurements were obtained from two-dimensional phase-contrast magnetic resonance imaging (2D PC MRI). After sacrifice at 180 days, the explants were collected for microstructural analysis. Histological sections were stained with Movat-Pentachrome and imaged using an Olympus Slideview VS200 slide scanner (Olympus, Waltham, MA, USA), equipped with a VS-264C camera.

ECM and scaffold volume fractions were extracted from these images with an in-house *Matlab R2021b* script. Pixels representing collagen and scaffold were identified by color thresholding, and each fraction was computed as the corresponding count divided by the sum of both. A similar approach is illustrated in Fig. 3 of our previous work (Shirazi et al., 2025). Blood pressures were measured through an arterial catheter in the right ear during the implantation and explantation surgeries, recording systolic, diastolic, and mean values.

### 2.2 Computational model

The computational model comprises two variants, which we denote FSG1 and FSG2, shown in Fig. 1. Both build on a finite element growth and remodeling framework that predicts the evolving graft properties, as detailed in Sections 2.2.1, 2.2.2, and 2.2.3. The framework updates these properties at every time step of one day and takes the time-averaged wall shear stress (TAWSS) as input. In the FSG1 model, the TAWSS follows from an artificial neural network (ANN) surrogate model trained on the results of 20 computational fluid dynamics (CFD) models of 20 different geometries (Section 2.2.4) and we describe the model and the ANN training in Section 2.2.6. In the FSG2 model, the TAWSS is updated every 60 time steps from a fluid-structure-interaction (FSI) model (Section 2.2.5), as detailed in Section 2.2.7.

**Figure 1:**
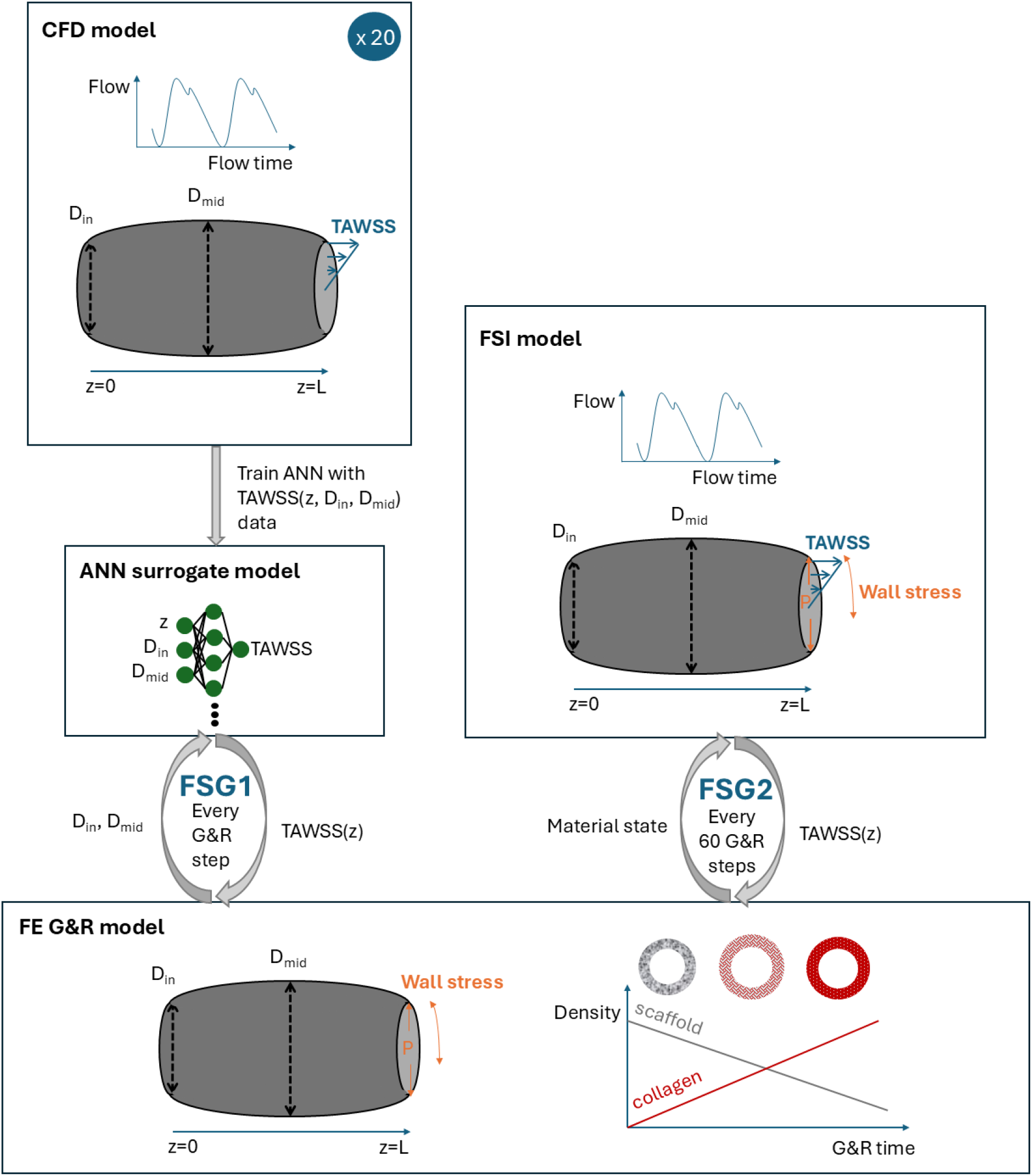
Overview of the proposed models. A finite element growth and remodeling (FE G&R) framework predicts the evolving graft properties from the time-averaged wall shear stress (TAWSS). In the FSG1 model, this TAWSS follows from an artificial neural network (ANN) surrogate model trained on 20 computational fluid dynamics (CFD) models of 20 different geometries, whereas in the FSG2 model it is updated every 60 time steps from a two-way coupled fluid-structure-interaction (FSI) model.

#### 2.2.1 Constitutive model

The graft wall is modeled as a hyperelastic material whose strain energy decomposes into contributions from the biodegradable scaffold (S), extracellular matrix collagen fibers (C) in four directions, and an isotropic ground matrix (GM) (Shirazi et al., 2025). Following Terzano et al. (2023), the strain-energy function of the scaffold reads

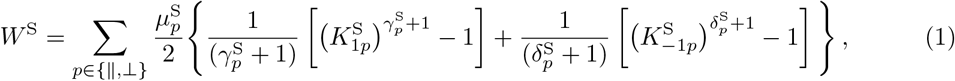

where 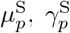 and 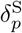 denote material parameters, and the invariants follows as 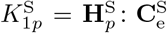 and 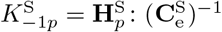, with *p* ∈ {∥, ⊥}. Here, 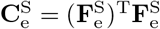 denotes the elastic right Cauchy–Green tensor of the scaffold, 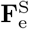 its elastic deformation gradient tensor, and 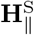 and 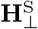 the coaxial and perpendicular structure tensors, respectively. A detailed derivation is given in Wollner et al. (2023).

The ground matrix is modeled as a compressible neo-Hookean material with strain-energy function given by (Shirazi et al., 2025)

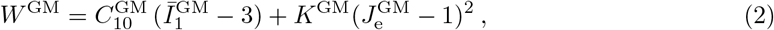

where 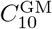 and 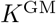 denote material parameters, and 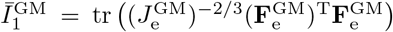 is the first invariant of the isochoric part of the right Cauchy–Green tensor, with 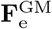 the elastic deformation gradient tensor of the ground matrix and 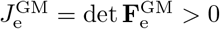.

In addition, we assume four collagen fiber families (*j* = 1, 2, 3, 4) with strain-energy function (Shirazi et al., 2025)

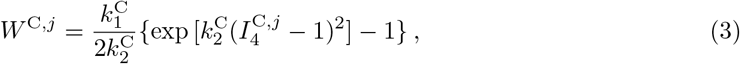

where 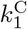 and 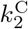 denote material parameters and 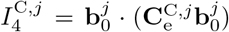 is the fourth invariant, with 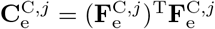 and 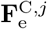 the elastic deformation gradient tensor of collagen fiber family *j*. The unit vector 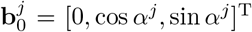 defines the reference fiber orientation in a cylindrical coordinate system whose first, second, and third directions are the radial, circumferential, and axial ones, with *α*^1^ = 0, *α*^2^ = *π/*2, and *α*^3^ = − *α*^4^ = *π/*4.

The total strain-energy function then follows as the weighted sum over all constituents (Shirazi et al., 2025)

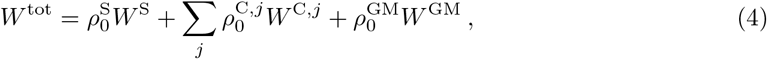

where 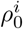 represents the mass density of constituent *i* at a certain time point per unit reference volume. The total Cauchy stress tensor of the mixture then reads

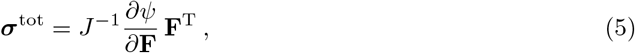

with **F** the total deformation gradient tensor of the mixture and *J* = det **F** *>* 0.

Following the homogenized constrained mixture theory (Cyron et al., 2016; Braeu et al., 2017) and assuming no inelastic volumetric growth, **F**_*g*_ = **I**, the elastic deformation gradient of the scaffold is given by

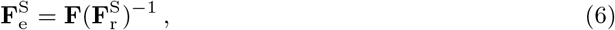

while for the ground matrix (*i* = GM) and collagen (*i* = C) we define

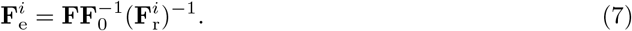

where **F**_0_ is the deformation gradient tensor at the initial time point, before extracellular matrix ingrowth, and 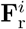 the inelastic remodeling deformation gradient of constituent *i*. In the reference configuration, we initialize 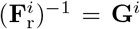, with **G**^*i*^ the deposition stretch tensor of constituent *i*, which, for the scaffold, ground matrix, and collagen, reads (Shirazi et al., 2025)

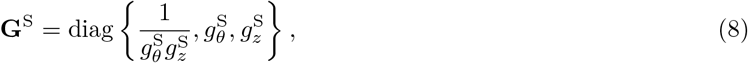

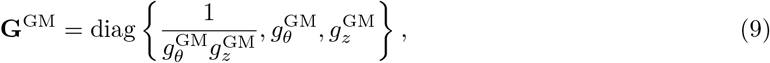

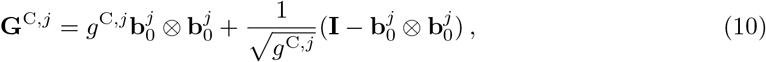

with 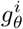 and 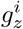 the deposition stretches of constituent *i* in the circumferential and axial directions, respectively, and *g*^C^ the deposition stretch of collagen along the fiber direction. The remodeling deformation gradient tensor 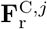 evolves over time following Maes and Famaey (2023), whereas 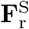 and 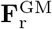 remain constant and equal to the inverse of the respective deposition stretch tensors.

#### 2.2.2 Growth and remodeling model

The growth and remodeling model is defined according to the homogenized constrained mixture theory (Cyron et al., 2016; Braeu et al., 2017), and accounts for production and degradation of the ECM (collagen and ground matrix) as well as for scaffold degradation. The scaffold material degrades as (Shirazi et al., 2025)

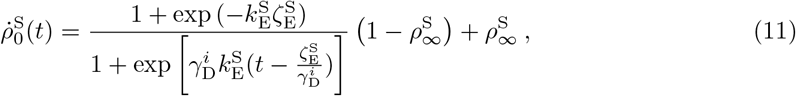

where 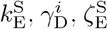, and 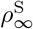 denote polymer degradation parameters and *t* the growth and remodeling time. The collage production rate is defined as (Shirazi et al., 2025)

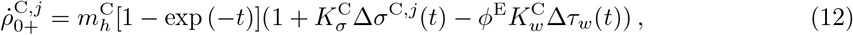

where 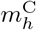 is the homeostatic collagen production rate and 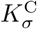 and 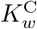 are gain parameters that scale production with deviation from the homeostatic fiber stress, 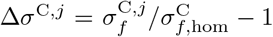, and from the homeostatic wall shear stress, Δ*τ*_*w*_ = *τ*_*w*_*/τ*_*w*,hom_ *−* 1. For each collagen fiber family, the fiber Cauchy stress follows as 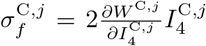, and the homeostatic stress 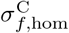 follows from the same expression evaluated at 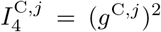. Here, *ϕ*^E^ denotes the local fraction of intact endothelium, which is required to sense the TAWSS. Sections 2.2.6 and 2.2.7 explain how we calculate *τ*_*w*_ and *τ*_*w*,hom_.

The collagen degradation rate reads (Shirazi et al., 2025)

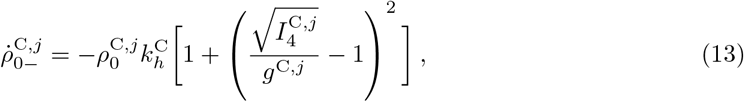

with the rate parameter 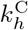, so that degradation accelerates as the current fiber stretch 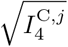 deviates from the deposition stretch *g*^C^. The ground matrix density follows the collagen density at all times through 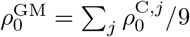 (90% collagen, 10% ground matrix).

Assuming smooth and continuous endothelial ingrowth from the anastomosis sites, we calculate *ϕ*^E^ as

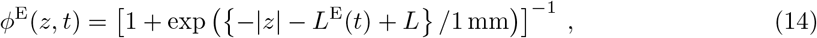

with the total conduit length *L*, axial position *z* ∈ [*L/*2, *L/*2], the time-dependent ingrowth length *L*^E^(*t*) = *Lt/*(2*T* ^E^), and *T* ^E^ the time until full endothelium ingrowth.

#### 2.2.3 Finite element model

We implemented the constitutive equations from Section 2.2.1 in a user-defined material (VUMAT) subroutine for explicit simulations in the commercial finite element software *Abaqus/Explicit* for the FSI models (see section 2.2.5). These equations, together with those from Section 2.2.2, were also implemented in a user-defined material (UMAT) subroutine for implicit simulations in *Abaqus/Standard* for the FSG models (see sections 2.2.6 and 2.2.7). Note the implementation of Eq. (1) was made available in previous work by Terzano et al. (2023), and Eqs. (1) and (3) by Maes and Famaey (2023). In addition, Eqs.(12) and (13) were implemented with a forward Euler integration scheme using time steps of one day, again following Maes and Famaey (2023).

Assuming axisymmetry, we defined the initial graft geometry as a sector of a hollow cylinder of length *L*, inner diameter *D*, and thickness *H*. The hexahedral mesh contains one element in circumferential direction, six in radial direction, and *n*_axial_ in axial direction. To allow the conduit wall to thicken by up to 50%, we added three additional elements as outer layer with an initial density 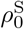 of 5% of that of the remaining wall, which has a negligible effect on the initial configuration.

In the initial simulation step, we apply an intraluminal pressure *p* and keep it constant thereafter. The inlet outlet boundaries are fixed in axial direction, no nodal displacements are allowed in circumferential direction, and radial displacement remains unconstrained. After initialization, we perform 180 or 360 growth and remodeling steps of one day and predict the evolution of the diameter and the changes in graft composition. Time and mesh convergence analyses were performed to ensure that discretization errors remain small.

We consider two case studies for which a comparison with experimental data is possible, the carotid conduit graft and the pulmonary valved conduit graft. Calibration data for both are available from literature (Brugmans et al., 2019; Shirazi et al., 2025), whereas we validate against the available experimental data presented in Section 2.1, in each case for two different initial graft thicknesses. This gives a total of six models, whose geometry and intraluminal pressure are detailed in Table 1.

**Table 1:** Geometrical parameters (*L*: length, *D*: inner diameter, and *H*: thickness) and intraluminal pressure *p* for both case studies, each comprising one calibration and two validation models with a thick or a thin wall. *n*_axial_ is the number of finite elements in axial direction.

| Parameter | Unit | Calibration |  | Validation |  |
| --- | --- | --- | --- | --- | --- |
|  |  | Carotid<br>(Shirazi et al., 2025) | Pulmonary<br>(Brugmans et al., 2019) | Carotid<br>(Section 2.1) | Pulmonary<br>(Section 2.1) |
| $L$ | (mm) | 45.0 | 35.0 | 70.0 | 27.0 |
| $D$ | (mm) | 6.0 | 20.8 | 6.0 | 20.0 |
| $H$ | (mm) | 0.50 | 1.0 | 0.63 (thick)<br>0.40 (thin) | 0.70 (thick)<br>0.57 (thin) |
| $p$ | (mmHg) | 80 | 10 | 78 | 10 |
| $n_{\text{axial}}$ | (–) | 112 | 140 | 175 | 68 |

#### 2.2.4 Computational fluid dynamics model

We define the CFD model of the conduit in *FlowVision* (Medividia Belgium) and prescribe a cyclic time-dependent flow rate at the inlet of the cylinder, as shown in Fig. 2. The flow depends on the case study and on whether the calibration or validation dataset is considered; see also Table 1. We assume rigid wall with no-slip condition and a zero-pressure free outlet, and we model blood as a Newtonian fluid with laminar flow. We simulate two cardiac cycles and evaluate the TAWSS at every surface node by time-averaging the wall shear stress over the second cycle, so that the flow is fully developed.

**Figure 2:**
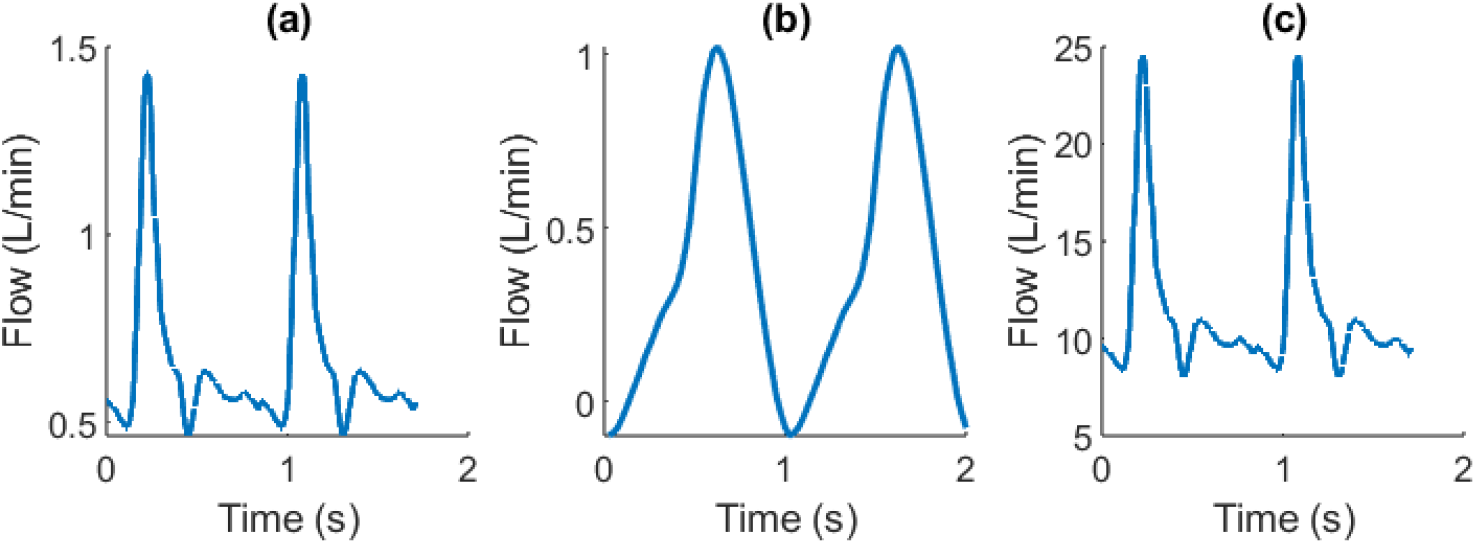
Imposed flow rates for (a) the carotid calibration, (b) the carotid validation, and (c) the pulmonary calibration and validation. The data in (a) were taken from Soleimani et al. (2017) and scaled to pulmonary artery level for (c), whereas the data in (b) were derived from 2D PC MRI as described in Section 2.1.

#### 2.2.5 Fluid-structure-interaction model

For the FSI model, we couple an *Abaqus/Explicit* finite element solver to the *FlowVision* computational fluid dynamics solver through the *Simulia* co-simulation engine. In Abaqus, we create a dummy surface part, whose outer surface nodes are tied to the inner surface of the solid geometry, so that the flow solver transfers the pressure loads through this tie constraint. The conduit is modeled as a solid part with three elements in radial direction and the material model is defined through the VUMAT implementation (cf. Section 2.2.3). At the start of the FSI simulation, we pressurize the structure, after which the fluid load smoothly takes over pressurization.

We again model blood as a Newtonian fluid. At the inlet, we prescribe the flow velocity and the pressure separately, using the same cyclic flow rate as in Section 2.2.4, and we fix the inlet and outlet pressure at the same level. As before, the TAWSS is evaluated at every surface node by time-averaging the wall shear stress over the second cardiac cycle.

#### 2.2.6 Fluid-structure-growth model – Version 1 (FSG1)

In the first version of the FSG model (FSG1), the TAWSS is calculated with an ANN surrogate model of the CFD model. To construct this surrogate model, we run 20 CFD models with 20 different geometries, all geometries were axisymmetric and symmetric with respect to the cross-sectional plane at mid-length. Twenty combinations of inlet, outlet (*D*_in_ = *D*_out_), and middle (*D*_mid_) diameters were obtained through Latin hypercube sampling with the *lhsdesign* function in *Matlab R2022b*. From these 20 samples of two geometrical parameters, we created smooth parts using the spline functionality in *Abaqus*. All 20 CFD models were run as explained in Section 2.2.4, after which the TAWSS was exported at the surface nodes.

Assuming axisymmetry of the results, the TAWSS depends on three parameters, namely *D*_in_, *D*_mid_, and the axial coordinate *z*. Because of the plug flow assumption at the inlet, the TAWSS is unreliable near the inlet and is therefore disregarded over the first 5% of the conduit length. An ANN was then trained with three input nodes (*D*_in_, *D*_mid_, and *z*), one fully connected layer of 10 nodes with sigmoidal activation, and one output node corresponding to the TAWSS. This architecture contains a limited amount of parameters to avoid overfitting of the noisy data. We trained the model with the *fitrnet* function in *Matlab R2022b*, using the LBFGS solver minimizing the mean squared error (MSE) through 1,000 iterations. The data, consisting of 500,000 to 1,700,000 points depending on the graft geometries, was standardized, and 80% of the data points was used for training and 20% for testing. The results of the training, with final losses between 0.06 and 4.41 Pa^2^ in terms of MSE, are shown in the supplementary material. Four different ANNs were trained, each corresponding to one of the two case studies (carotid or pulmonary) and either for model calibration or model validation. These four ANNs hence correspond to the four cases detailed in Table 1.

Given *D*_in_ and *D*_mid_ at any growth and remodeling time point, the resulting ANN yields the TAWSS at every axial location *z*. The TAWSS, denoted *τ*_*w*_, then fully defines the collagen production rate at the same time and location through Eq. (12). In practice, this ANN was implemented as a *Fortran* subroutine called by the UMAT. At every time point, *D*_in_ and *D*_mid_ were determined through a *URDFIL* utility subroutine for *Abaqus*, using *DBFILE* to access the nodal coordinates after every increment. The homeostatic TAWSS, denoted *τ*_*w*,hom_, is calculated at baseline, that takes the initial graft diameter for *D*_in_ and *D*_mid_ into consideration.

#### 2.2.7 Fluid-structure-growth model – Version 2 (FSG2)

In the second version of the FSG model, the TAWSS is calculated directly from the FSI model explained in Section 2.2.5. A first FSI model is run at baseline conditions, that is, with only scaffold material present and with the initial graft dimensions right after implantation. The resulting TAWSS is then exported to the ETR model by defining at every node of the solid ETR model a TAWSS field variable that takes values of the closest surface node of the fluid model. Here again, the TAWSS is ignored for the first 5% of the conduit length. These baseline values (*τ*_*w*,hom_) are defined as the homeostatic TAWSS and also serve as current TAWSS in the first 60 ETR simulation steps. After these 60 steps, another FSI model is run with updated localized material properties, after which the current TAWSS values (*τ*_*w*_) are again imported into the ETR model. This process is repeated until the intended total growth and remodeling time is reached.

## 3 Results

### 3.1 Model calibration

The scaffold degradation parameters 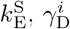, and 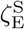 from Eq. (11) were calibrated first by fitting scaffold density data obtained from histology (Brugmans et al., 2019; Shirazi et al., 2025). The fit was performed with *lsqnonlin* function in *Matlab R2021b*, minimizing the objective function

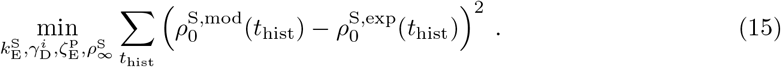

Here, *t*_hist_ denotes the time points at which histology data is available, 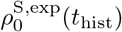 the experimentally measured scaffold density at those times, and 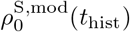 the corresponding density predicted by the model through Eq. (11). In the pulmonary conduit case, the additional parameter 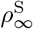 was optimized as well, whereas it was assumed to be zero in the carotid conduit case.

To calibrate the remaining model parameters, a semi-analytical model similar to the one reported in Shirazi et al. (2025) was implemented in *Matlab R2021b*. This simplified version of the FSG1 model from Section 2.2.6 facilitates calibration of the model parameters. The implementation includes the ANN as surrogate model of the CFD model, which captures TAWSS-dependent remodeling through Eq. (12). Two separate models, one for the inlet and one for the mid-conduit location, were run simultaneously so that the TAWSS could be calculated of the TAWSS with the network. Using the *lsqnonlin* function in *Matlab R2021b*, this model was run with several guesses of the parameters 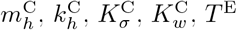, and 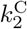 to minimize the following objective function

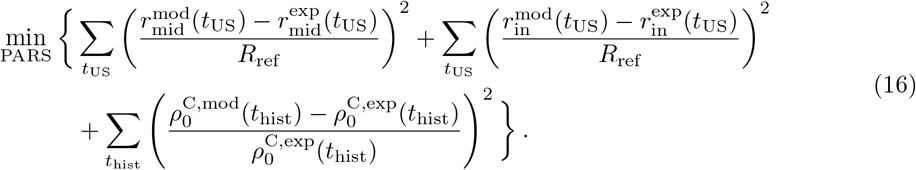

Here, *t*_US_ denotes the time points at which ultrasound data is available, proving the inner conduit radius at inlet and at mid-section, 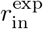 and 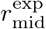, respectively. At those time points, the difference with the corresponding model-predicted inner radii, 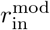 and 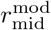, is minimized. The same applies for the collagen densities measured from histology and predicted by the model though Eq. (12), denoted 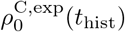 and 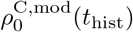, respectively. The parameter *R*_ref_ acts as a scaling factor.

Where necessary, the collagen production rate parameters 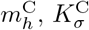, and 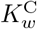 were further fine-tuned by trial and error to account for potential errors introduced by the simplified model so that the predictions of the FSG1 model (Section 2.2.6) matched the experimental data entering in Eq. (16) (Brugmans et al. (2019) for the pulmonary case study and Shirazi et al. (2025) for the carotid case study). The remaining parameters were fixed based on our previous study (Shirazi et al., 2025).

Table 2 gives an overview of all parameter values for both case studies, i.e., the carotid conduit and the pulmonary conduit. Figure 3 shows the calibration results for the carotid case study, where model predictions are compared to experimental measurements of radius and of scaffold and collagen densities. Similar results for the pulmonary case study are shown in Fig. 4.

**Table 2:** Overview of the parameter values for both case-studies: the carotid conduit and the pulmonary conduit. As mentioned in the last column, some parameters are obtained from literature, some from the model calibration activities explained in Section 3.1.

|  | Carotid | Pulmonary | Source |
| --- | --- | --- | --- |
| **Scaffold** |  |  |  |
| $H_{11}$ (-) | 0.691765 | 0.704960 | Terzano et al. (2023) |
| $H_{22}$ (-) | 0.258235 | 0.290040 | Terzano et al. (2023) |
| $\mu_{\parallel}^S$ (MPa) | 0.171399 | 1.533770 | Terzano et al. (2023) |
| $\mu_{\perp}^S$ (MPa) | 0.199557 | 0.802883 | Terzano et al. (2023) |
| $\gamma_{\parallel}^S$ (-) | 0.000699 | 0.000164 | Terzano et al. (2023) |
| $\gamma_{\perp}^S$ (-) | 0.291492 | 0.507081 | Terzano et al. (2023) |
| $\delta_{\parallel}^S$ (-) | 0.213387 | 0.586208 | Terzano et al. (2023) |
| $\delta_{\perp}^S$ (-) | 0.170464 | 0.358070 | Terzano et al. (2023) |
| $D_1$ (MPa$^{-1}$) | 0.548286 | 0.087043 | Terzano et al. (2023) |
| $g_{\theta}^S$ (-) | 1.01 | 1.01 | Shirazi et al. (2025) |
| $g_z^S$ (-) | 1.01 | 1.01 | Shirazi et al. (2025) |
| **Collagen** |  |  |  |
| $k_1^C$ (MPa) | 0.010 | 0.010 | Shirazi et al. (2025) |
| $k_2^C$ (-) | 127.8 | 38.23 | Section 3.1 |
| $g^C$ (-) | 1.08 | 1.08 | Shirazi et al. (2025) |
| **Ground matrix** |  |  |  |
| $C_{10}^{GM}$ (MPa) | 0.050 | 0.050 | Shirazi et al. (2025) |
| $K^{GM}$ (MPa) | 0.250 | 0.250 | Shirazi et al. (2025) |
| $g_{\theta}^{GM}$ (-) | 1.1664 | 1.1664 | Shirazi et al. (2025) |
| $g_z^{GM}$ (-) | 1.1664 | 1.1664 | Shirazi et al. (2025) |
| **G&R** |  |  |  |
| $k_E^S$ (days) | 0.0270 | 0.1721 | Section 3.1 |
| $\gamma_D^i$ (-) | 0.3 | 0.3 | Section 3.1 |
| $\zeta_E^S$ (days) | 5.0 | 0.1 | Section 3.1 |
| $\rho_{\infty}^S$ (-) | 0.0 | 0.3449 | Section 3.1 |
| $m_h^C$ (days$^{-1}$) | 0.0182 | 0.0043 | Section 3.1 |
| $K_{\sigma}^C$ (-) | 0.0100 | 0.470 | Section 3.1 |
| $K_w^C$ (-) | 2.93 | 0.489 | Section 3.1 |
| $k_h^C$ (days$^{-1}$) | 0.0611 | 0.0254 | Section 3.1 |
| $T^E$ (days) | 35 | 35 | Section 3.1 |

**Figure 3:**
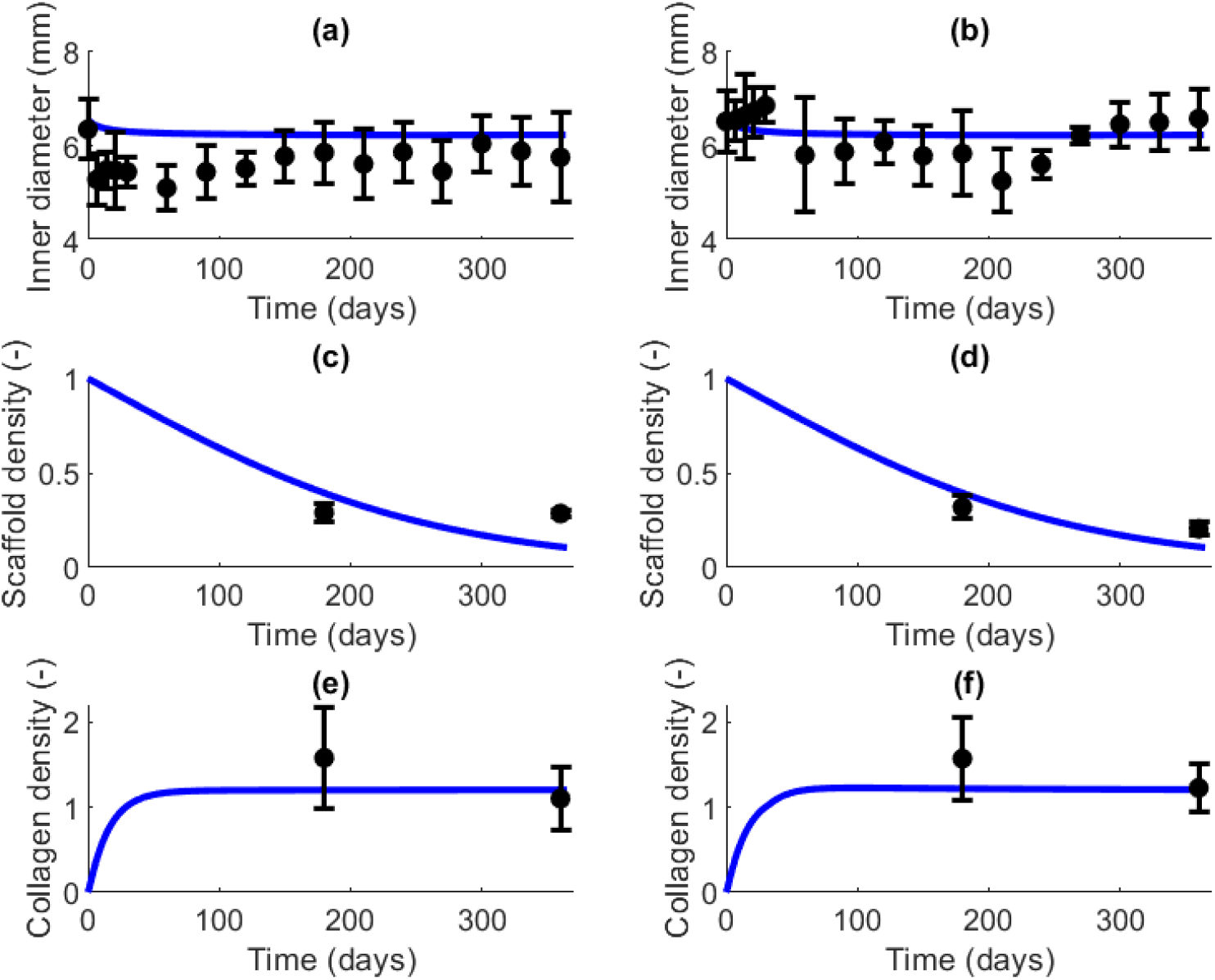
Results of the calibration of the FSG1 model for the carotid case study. Model predictions (solid curves) are compared to experimentally measured means and standard deviations (error bars) from Shirazi et al. (2025). (a,b) Inner diameter, (c,d) scaffold density, and (e,f) collagen density. The left column (a,c,e) shows the results for the inlet location, the right column (b,d,f) those at the mid-graft location.

**Figure 4:**
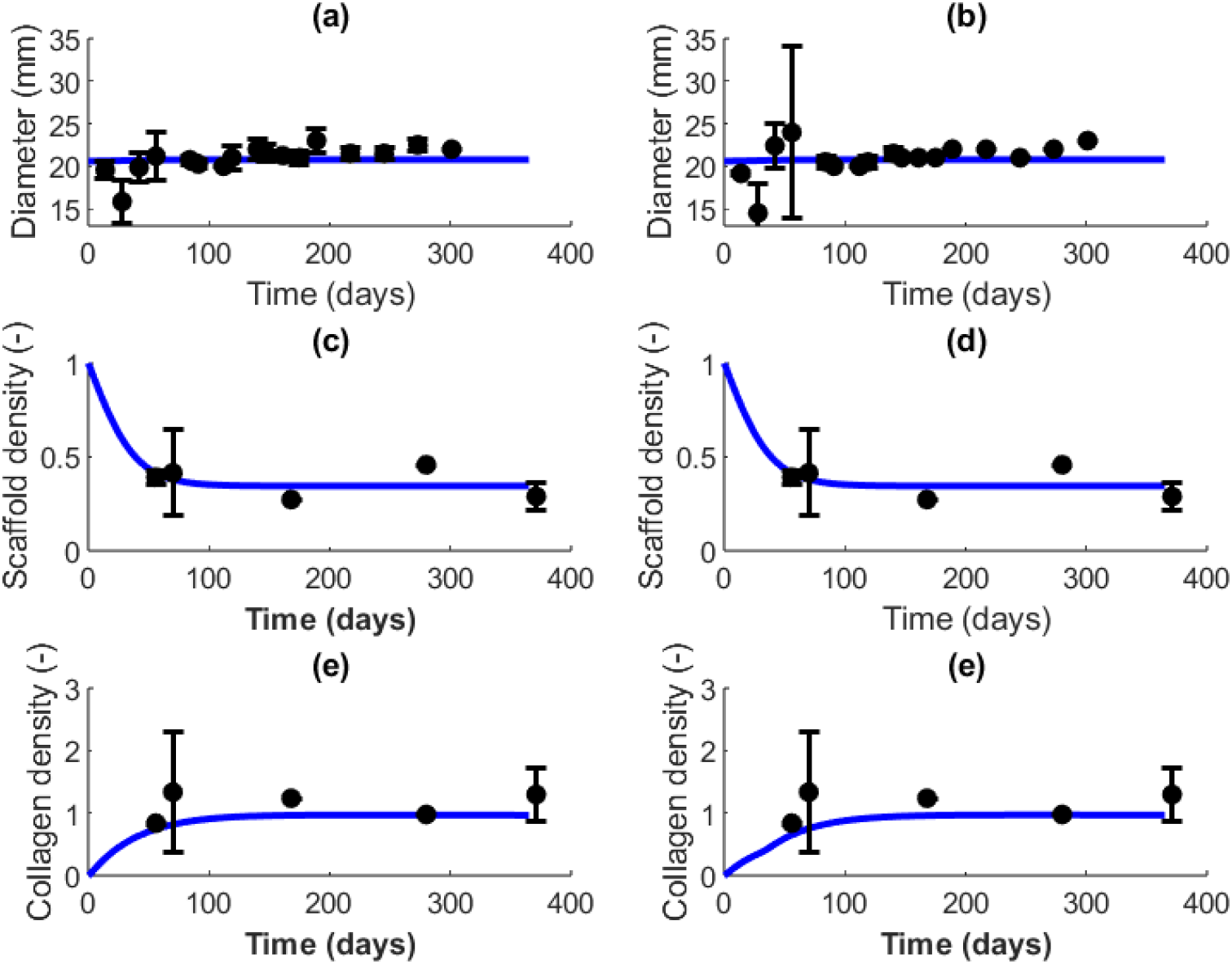
Results of the calibration of the FSG1 model for the pulmonary case study. Model predictions (solid curves) are compared to experimentally measured means and standard deviations (error bars) from Brugmans et al. (2019). (a,b) Inner diameter, (c,d) scaffold density, and (e,f) collagen density. The left column (a,c,e) shows the results at the inlet location, the right column (b,d,f) at the mid-graft location.

### 3.2 Model verification

#### 3.2.1 Comparisons of TAWSS predictions across models

The FSG1 model relies on an ANN as surrogate model of the CFD model, which in turn replaces the computationally expensive FSI model. The first verification activity therefore quantifies the error introduced when the TAWSS is calculated with the ANN rather than with the CFD or FSI model. For the carotid case study, results are shown in Fig. 5 for *D*_mid_ = 4.3 mm and *D*_in_ = *D*_out_ = 7.1 mm. Qualitatively, the three models agree well. Using the *goodnessOfFit Matlab* function, the normalized root mean square error (NRMSE) of ANN-predicted TAWSS as a function of the axial coordinate *z* amounts to 12.8% with respect to the CFD result and 12.6% with respect to the FSI result.

**Figure 5:**
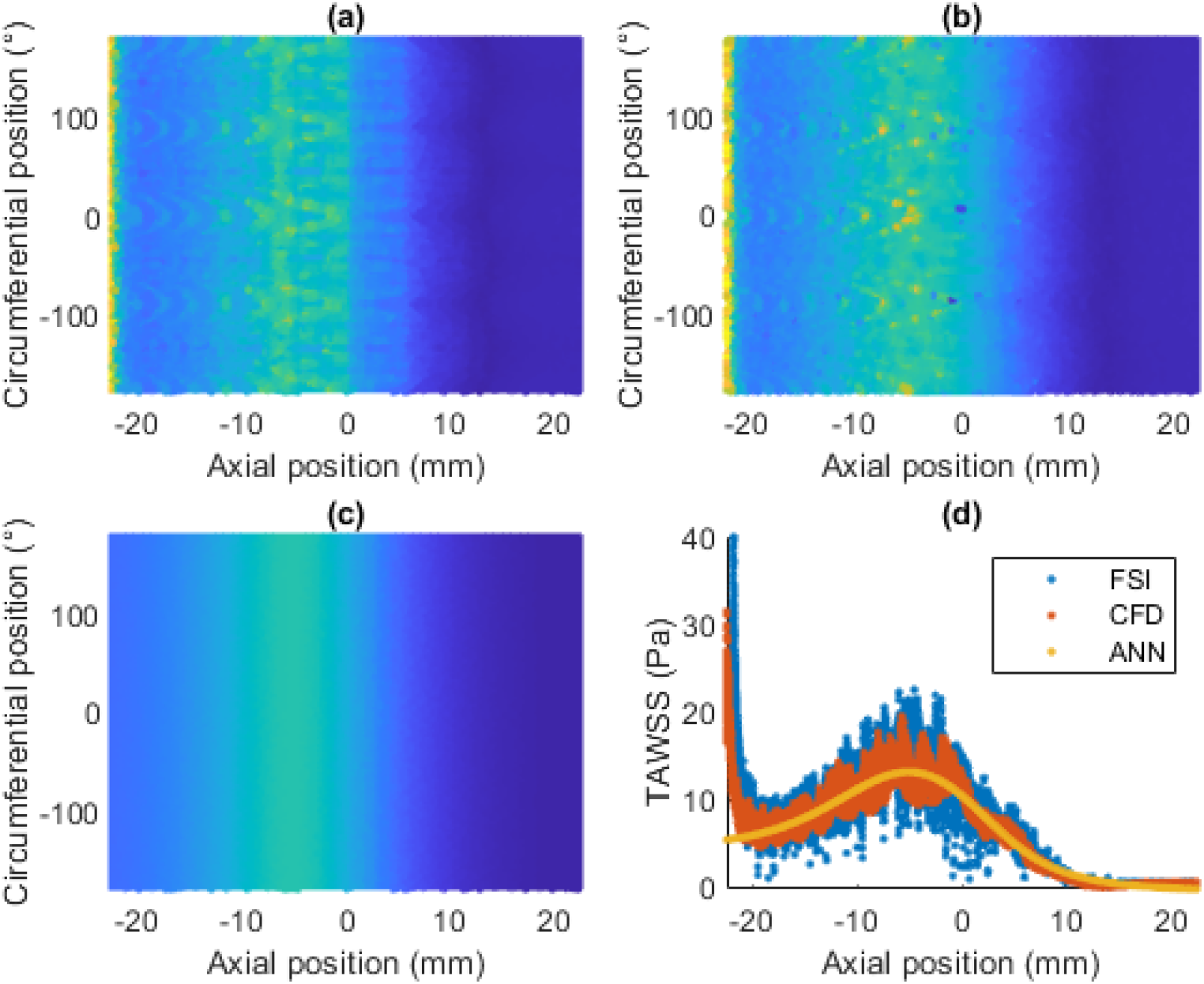
Comparison of time-averaged wall shear stress (TAWSS) predictions from the computational fluid dynamics (CFD), fluid-structure interaction (FSI), and surrogate models (artificial neural network (ANN)) for the carotid dataset of Shirazi et al. (2025). (a,b,c) Color maps of the predicted TAWSS as a function of axial and circumferential position, obtained with the CFD model, the FSI model, and the surrogate model, respectively. (d) Predicted TAWSS as a function of axial position for the three models.

#### 3.2.2 Comparison of growth and remodeling predictions between FSG1 and FSG2 models

The second verification activity compares the growth and remodeling prediction results of the FSG1 and FSG2 models. Both are also compared to an equivalent model that simply assumes

Hagen-Poiseuille flow, for which Eq. (12) uses 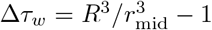, with *R*_mid_ and *r*_mid_ the initial and the current graft radius at mid-graft location, respectively. The three models were run with the geometry and the model parameters for the carotid case study. Figure 6 shows the evolving diameters, scaffold densities, and collagen densities of each model at the inlet and mid-graft locations over a prediction period of 180 days. The scaffold density results are identical, because its evolution depends on time only (cf. Eq. (11)) and is therefore not influenced by model-specific wall stress or TAWSS calculations. The relative difference in predicted collagen density between FSG1 and FSG2, on the other hand, is largest at the mid-graft location after 180 days of growth and remodeling, where it reaches 9.79%. This difference follows from the dependence of collagen production on the mechanical stimuli and on re-endothelialization (cf. Eq. (12)). The relative differences in predicted diameter remain small and stay below 1% over the prediction time frame.

**Figure 6:**
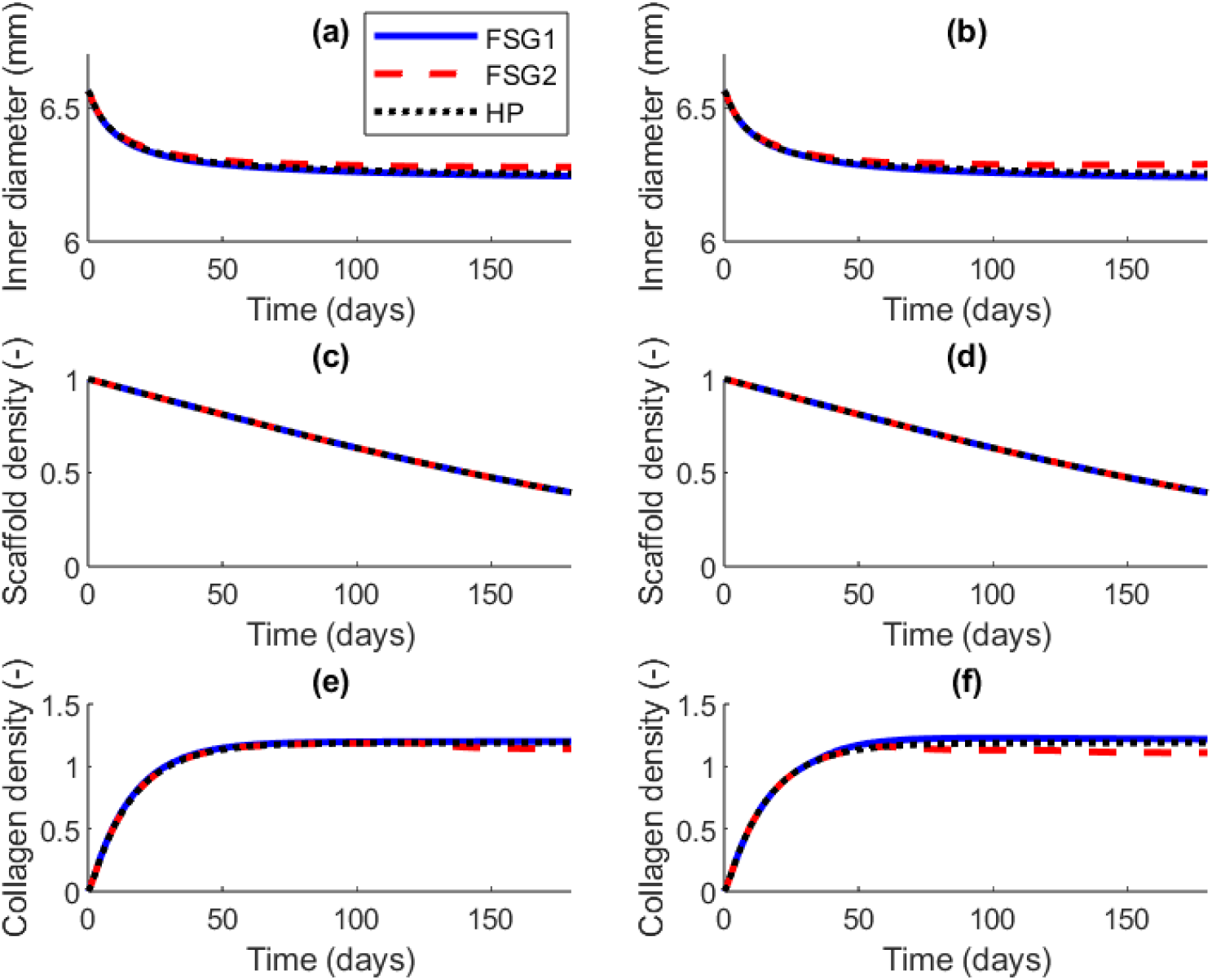
Comparison of growth and remodeling predictions from the FSG1 and FSG2 models for the carotid dataset of Shirazi et al. (2025). FSG1 model predictions (blue solid curves) are compared to FSG2 model predictions (red dashed lines) and to the predictions of the Hagen-Poiseuille (HP) model (black dotted lines). (a,b) Inner diameter, (c,d) scaffold density, and (e,f) collagen density. The left column (a,c,e) shows all results at the inlet location, the right column (b,d,f) those at the mid-graft location.

### 3.3 Model validation

For model validation, the FSG1 model is defined for both the carotid and the pulmonary conduit graft study using the experimental data generated in this work. The corresponding geometries and boundary conditions are listed in the last two columns of Table 1. For each case study, two models are run, one for the thick and one for the thin initial wall thickness of the implanted grafts.

Figure 7 shows how the predicted evolution of the diameter and the constituent fractions compares with the experimental measurements for the carotid case study. Model and experiment match well for the constituent fractions, defined as 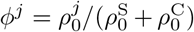 with *j* = C,S. The diameter is correctly predicted for the initial stage of growth and remodeling. After one or two months, however, the experiments show pronounced graft dilation that the model does not capture, because this effect is absent from the calibration data. Figure 8 shows the validation results for the pulmonary case study. Experiment and model predictions match well overall, in this case also over longer time periods.

**Figure 7:**
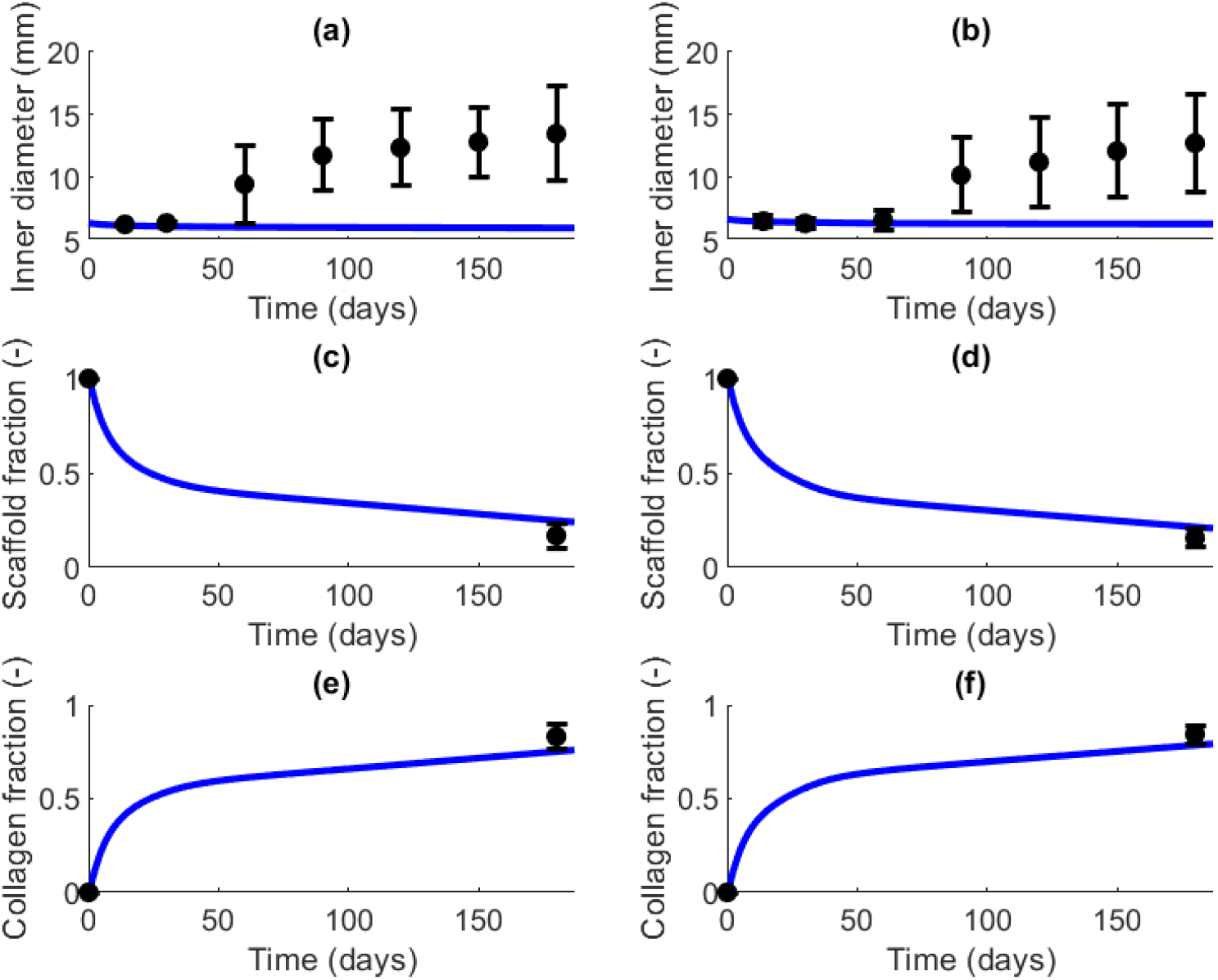
For the carotid case study, model predictions (solid curves) are compared to experimentally measured means and standard deviations (error bars) from the validation data of Shirazi et al. (2025). (a,b) Inner diameter, (c,d) scaffold fraction, and (g,f) collagen fraction. The left column (a,c,e) shows all results for the thick graft (inner wall thickness 0.63 mm), the right column (b,d,f) those for the thin graft (0.40 mm).

**Figure 8:**
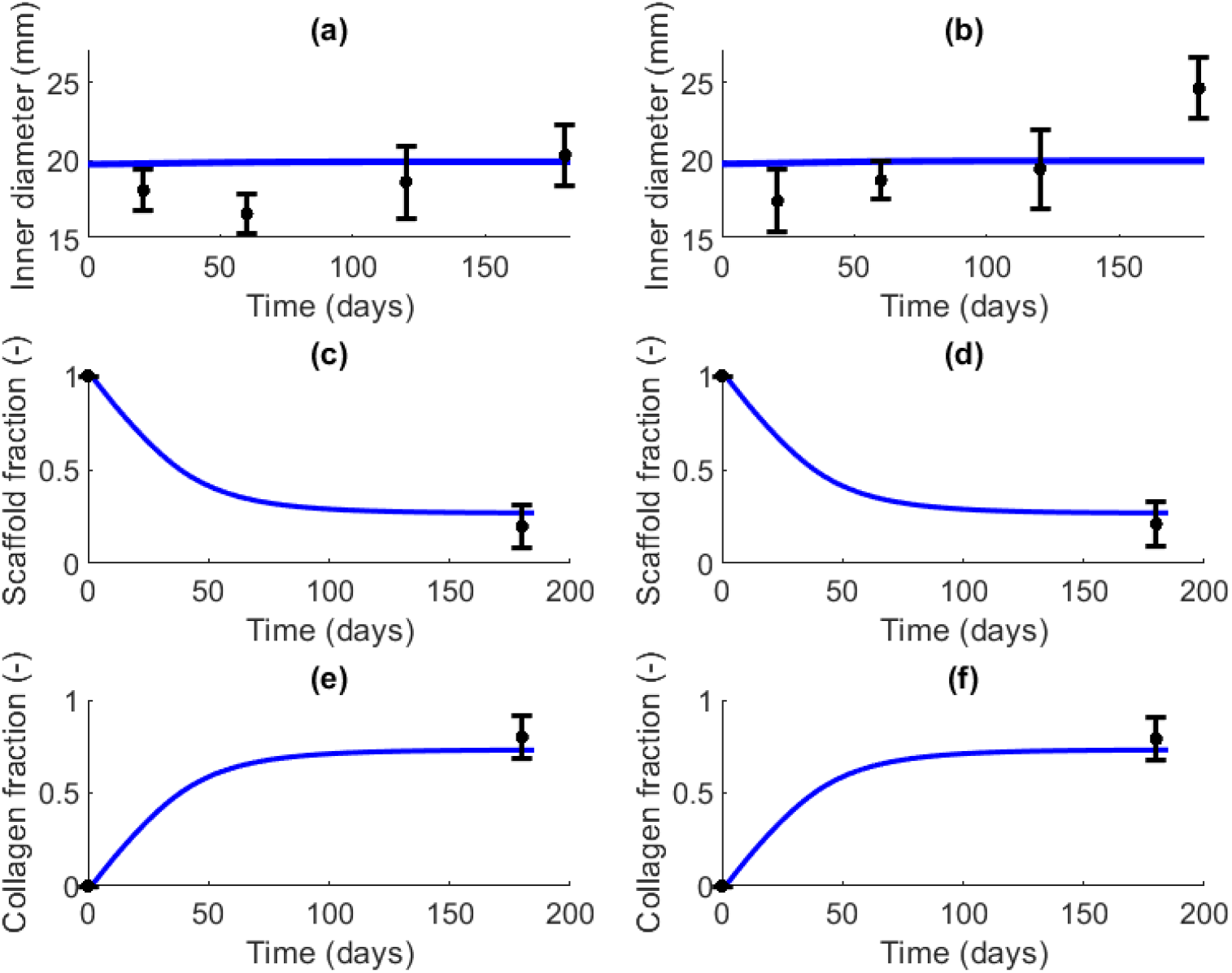
For the pulmonary case study, model predictions (solid curves) are compared to experimentally measured means and standard deviations (error bars) from the experimental validation data of Brugmans et al. (2019). (a,b) Inner diameter, (c,d) scaffold fraction, and (g,f) collagen fraction. The left column (a,c,e) shows all results for the thick graft (wall thickness 0.70 mm), the right column (b,d,f) those for the thin graft (0.57 mm).

## 4 Discussion

This study explores the feasibility and reliability of computationally efficient FSG models in the context of ETR. We propose two alternative approaches. In the first (FSG1), the TAWSS is calculated at every time point of growth and remodeling through an ANN surrogate model trained on CFD data. In the second (FSG2), the TAWSS is updated less frequently, but with a high-fidelity two-way coupled FSI model. Our verification study shows that predicted TAWSS is similar across the different calculation methods, i.e., the FSI model, the CFD model, and the ANN surrogate model. Moreover, we show that the FSG1 and FSG2 models predict similar time evolutions of graft diameter and constitution for the defined boundary value problem of a symmetric tube under inner pressure. This agreement is remarkable given the two very different model implementations, i.e., continuous updating of a low-fidelity TAWSS in FSG1 against occasional updating of a high-fidelity TAWSS in FSG2. We therefore conclude that a fully coupled FSG model with continuous update of TAWSS is likely unnecessary to reach acceptable results for the given constitutive model implementation. Figure 6 also shows that Hagen-Poiseuille assumption for the TAWSS performs similarly, which suggests that the model form uncertainty likely smaller than parameter uncertainty. The more strongly collagen production depends on th TAWSS through the parameter 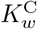, however, the more accurate the TAWSS prediction has to be, and the more the two models may deviate from each other and from the Hagen-Poiseuille solution. In contrast to our results, Schwarz et al. (2023) obtained different predictions with a fully coupled FSG solver than with the Hagen-Poiseuille equation. The present study assumes a simple conduit graft, whereas they also considered valve motion, for which the flow patterns likely deviate more from the idealized Hagen-Poiseuille flow.

Both FSG models are calibrated to two different applications, the carotid conduit and the pulmonary conduit. The carotid artery is part of the systemic circulation and therefore operates in a high pressure regime, whereas the pulmonary artery is the main artery of the pulmonary circulation and therefore operates in a lower pressure regime. Parameter values defining the mechanical properties of the scaffold material are taken from a previous study (Terzano et al., 2023), while growth and remodeling parameters are calibrated to match the measured time evolution of graft diameter and constitution as closely as possible. This calibration is needed because the growth and remodeling model of Shirazi et al. (2025) is adapted here to account for TAWSS-dependent collagen production. The model contains many phenomenological parameters, so that the solution found is potentially not unique and uncertainty may remain. To avoid returning a local optimum, multiple start points were tested during parameter optimization. Overall, Figs. 3 and 4 show a good match between model and experiment both case studies, although the spread on the experimental data is considerable. Probabilistic modeling techniques should be considered in future work to account for this uncertainty (Ranftl et al., 2026).

Considering also the results of the validation study (cf. Figs. 7 and 8), the model credibility analysis based on the experimental datasets of this study is convincing, except for the prediction of the diameter after two months in the carotid conduit case (cf. Fig. 7(a-b)). Since the experimentally observed graft dilatation is absent from the data used for model calibration (cf. Fig. 3(a-b)), the inability of the model to predict this behaviour is logical. It is currently unclear what causes these large differences given the similar experimental designs. The most important difference between both, as Table 1 shows, is the length of the implanted graft. Future experimental and computational work should therefore investigate how differences in graft length influence progressive graft dilatation. The present model introduces a length through the gradual ingrowth of the endothelium (cf. Eq. (14)), but the validation data show that the dependence is not sufficient. A further, unreported analysis shows that model does not predict the observed dilatation after recalibration on the validation data. This is achieved mainly through a reduction in the homeostatic collagen production rate 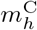, a reduction of the collagen strain-stiffening parameter 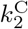, and an increase in 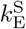 that accelerates scaffold degradation. Together, these three effects progressive increase graft compliance and thereby produce dilatation.

Before applying the proposed FSG1 model to other case studies, some limitations need to be addressed. The ANN used to calculate the TAWSS is trained on case-specific data, with the graft length, approximate diameter, the perfectly cylindrical shape, and the flow rate pattern all predefined. A new ANN therefore needs to be trained for each new application. Future work can consider more sophisticated surrogate models that take additional flow and geometry parameters as input variables. Moreover, we verified, calibrated, and validated our model on simple conduit geometries only. Especially in the context of ETR, more complex geometries and flow patterns, for example including valve leaflets and their motion, are necessary.

We conclude that the proposed FSG models are reliable in predicting the course of ETR in biodegradable vascular grafts, although caution is warranted when extrapolating to different case studies. Moreover, while short-term predictions are properly validated, the long-term predictions require more careful consideration.

## Data availability

The model files to reproduce the presented results are freely accessible at KU Leuven RDR through https://doi.org/10.48804/V4JQYQ (Maes et al., 2026).

