## Supplementary material for "Feasibility study of fluid-structure-growth modeling in the context of endogenous tissue restoration"

[a]BioMechanics, Department of Mechanical Engineering, KU Leuven, Leuven, Belgium
[b]Cardiac Surgery, Department of Cardiovascular Sciences, KU Leuven, Leuven, Belgium
[c]Capvidia NV, Leuven, Belgium
[d]Medividia BV, Herent, Belgium
[e]4RealSim Services BV, IJsselstein, The Netherlands
[f]Institute of Applied Mechanics, Friedrich-Alexander-Universität Erlangen-Nürnberg, Erlangen, Germany
[g]Institute of Biomechanics, Graz University of Technology, Graz, Austria
[h]MINES Saint-Etienne
[i]Xeltis BV, Eindhoven, The Netherlands
[j]Department of Structural Engineering, Norwegian University of Science and Technology (NTNU), Trondheim, Norway


### 1 Results of the surrogate model training

Figures 1 through 8 show the results of the artificial neural network (ANN) training as surrogate model for computational fluid dynamics (CFD) models. Specifically, for each of the two case-studies, carotid and pulmonary, two ANNs are trained for the calibration and validation datasets. For each of these in total four ANNs, training data was used with the axial (z) coordinate, inlet diameter ($D_{in}$), and mid-graft diameter ($D_{mid}$) as input parameters, and time-averaged wall shear stress as output parameters. The figures show how the ANN results match the CFD results.

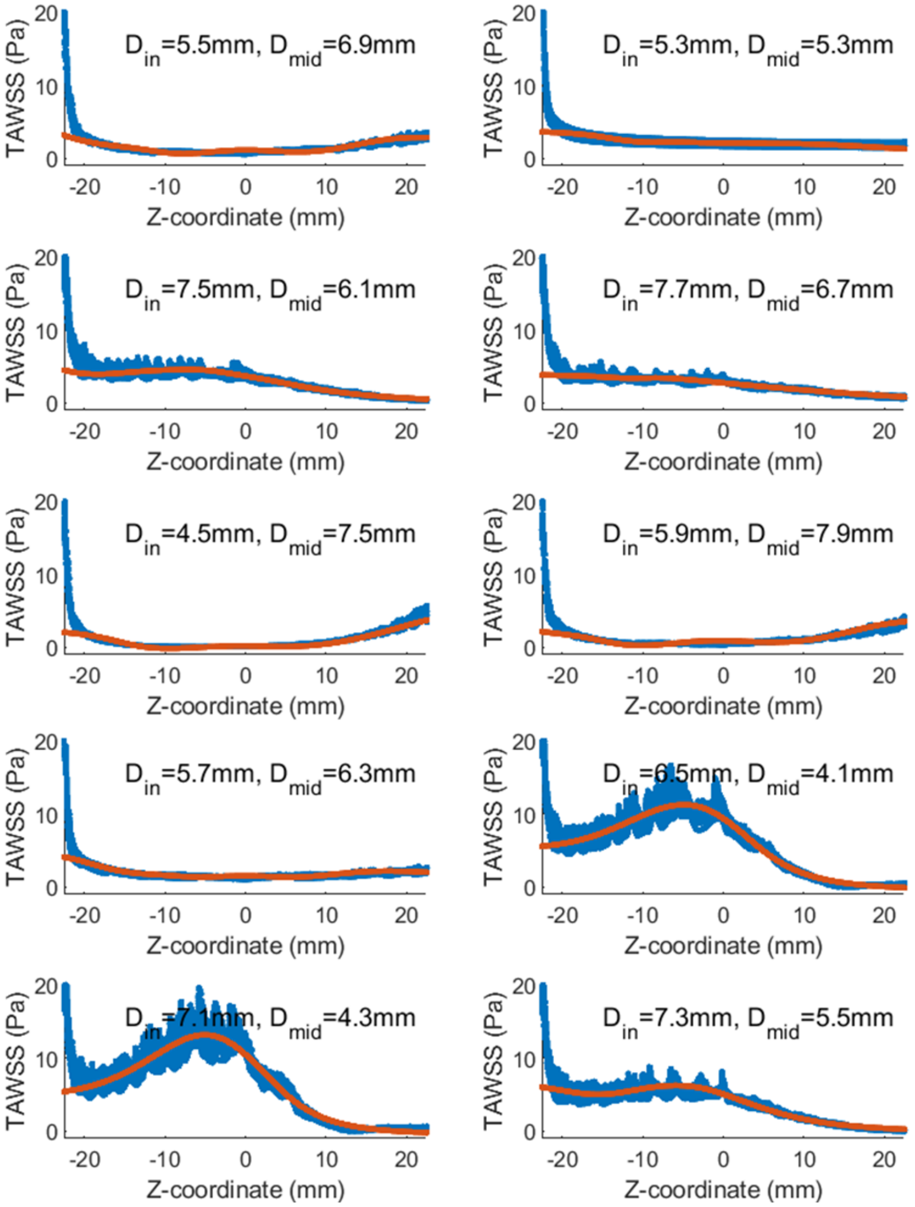


Figure 1: Results of the surrogate model training for the carotid calibration case. The TAWSS from the CFD simulation (blue) and the ANN (red) is shown in function of the Z-coordinate for the ten first combinations of $D_{in}$ and $D_{mid}$.

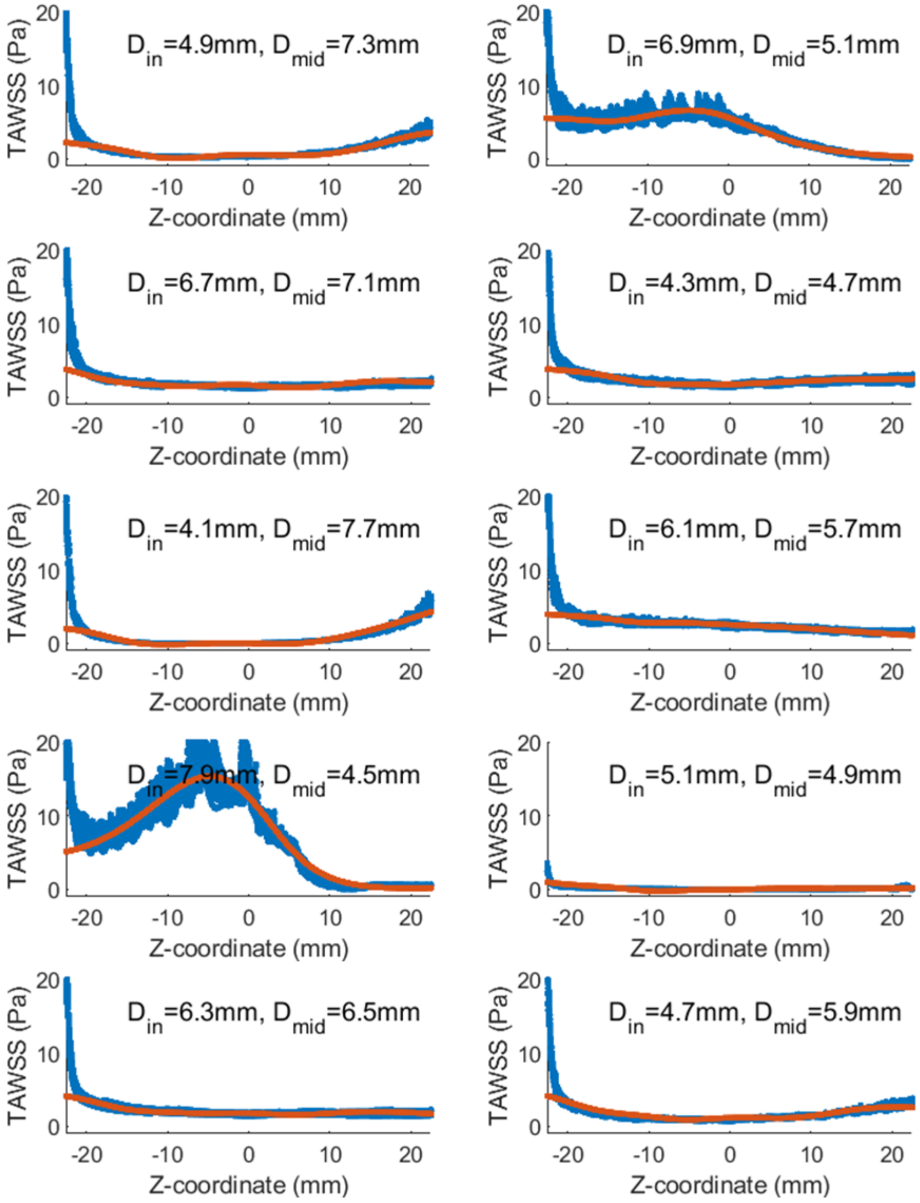


Figure 2: Results of the surrogate model training for the carotid calibration case. The TAWSS from the CFD simulation (blue) and the ANN (red) is shown in function of the Z-coordinate for the ten last combinations of $D_{in}$ and $D_{mid}$.

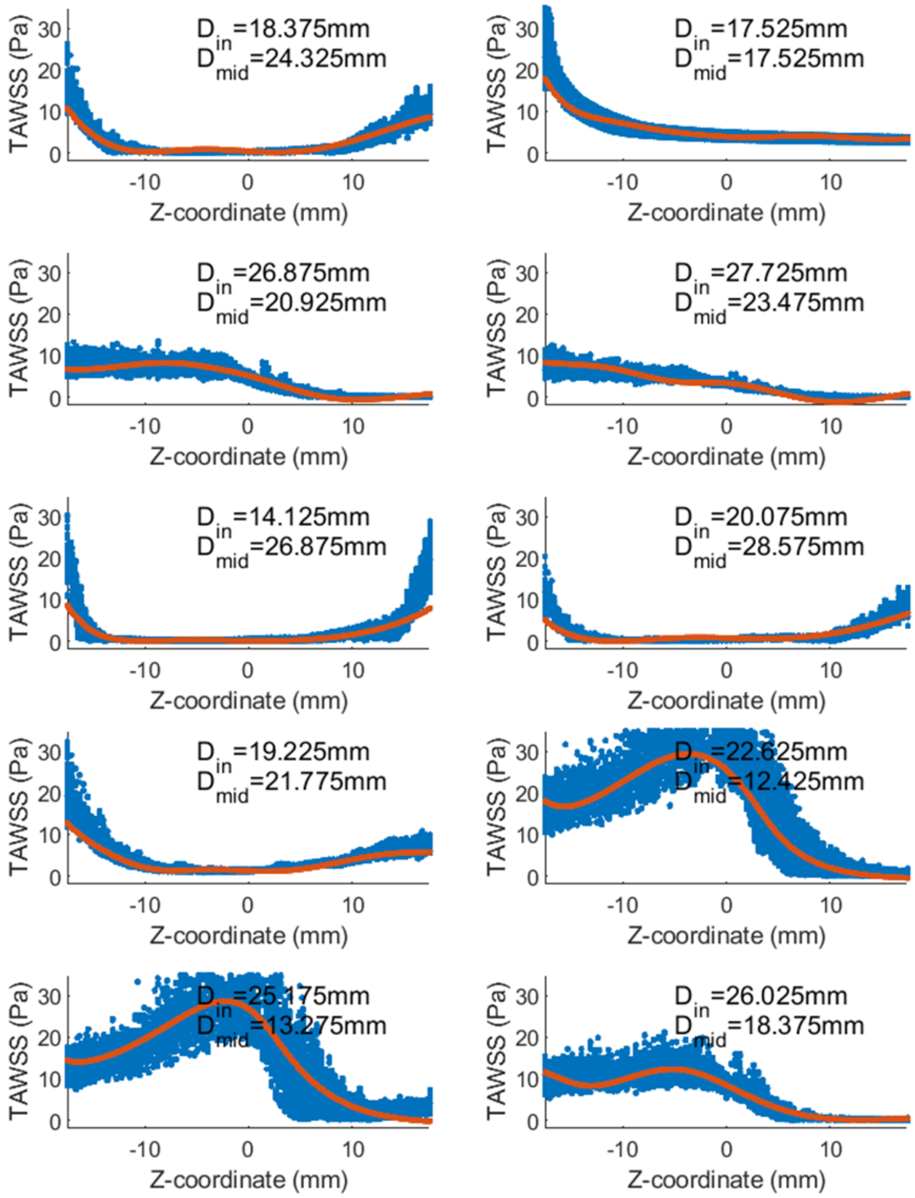


Figure 3: Results of the surrogate model training for the pulmonary calibration case. The TAWSS from the CFD simulation (blue) and the ANN (red) is shown in function of the Z-coordinate for the ten first combinations of $D_{in}$ and $D_{mid}$.

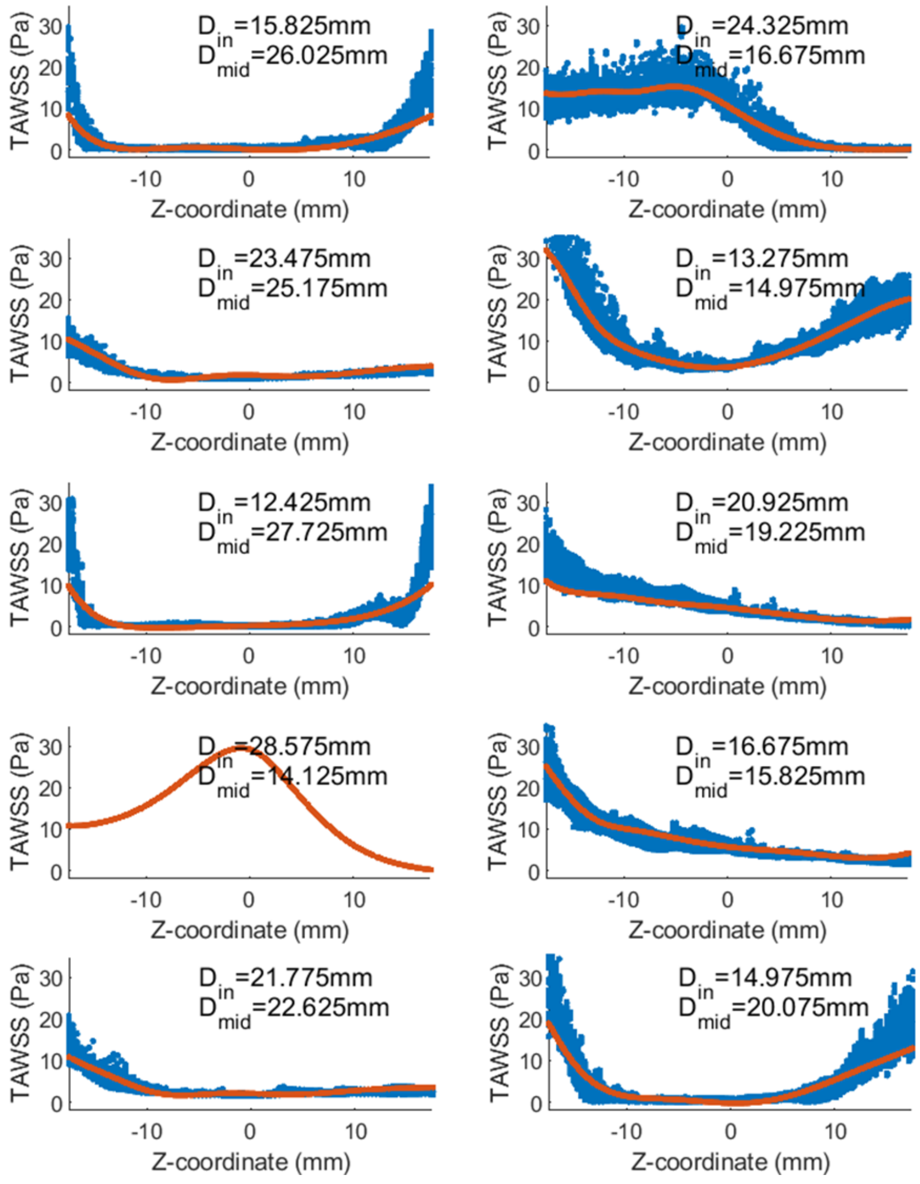


Figure 4: Results of the surrogate model training for the pulmonary calibration case. The TAWSS from the CFD simulation (blue) and the ANN (red) is shown in function of the Z-coordinate for the ten last combinations of $D_{in}$ and $D_{mid}$. No stable CFD solution could be found for the case with $D_{in}$=28.575mm and $D_{mid}$=14.125mm. This case was hence not taken into account for the training of the ANN.

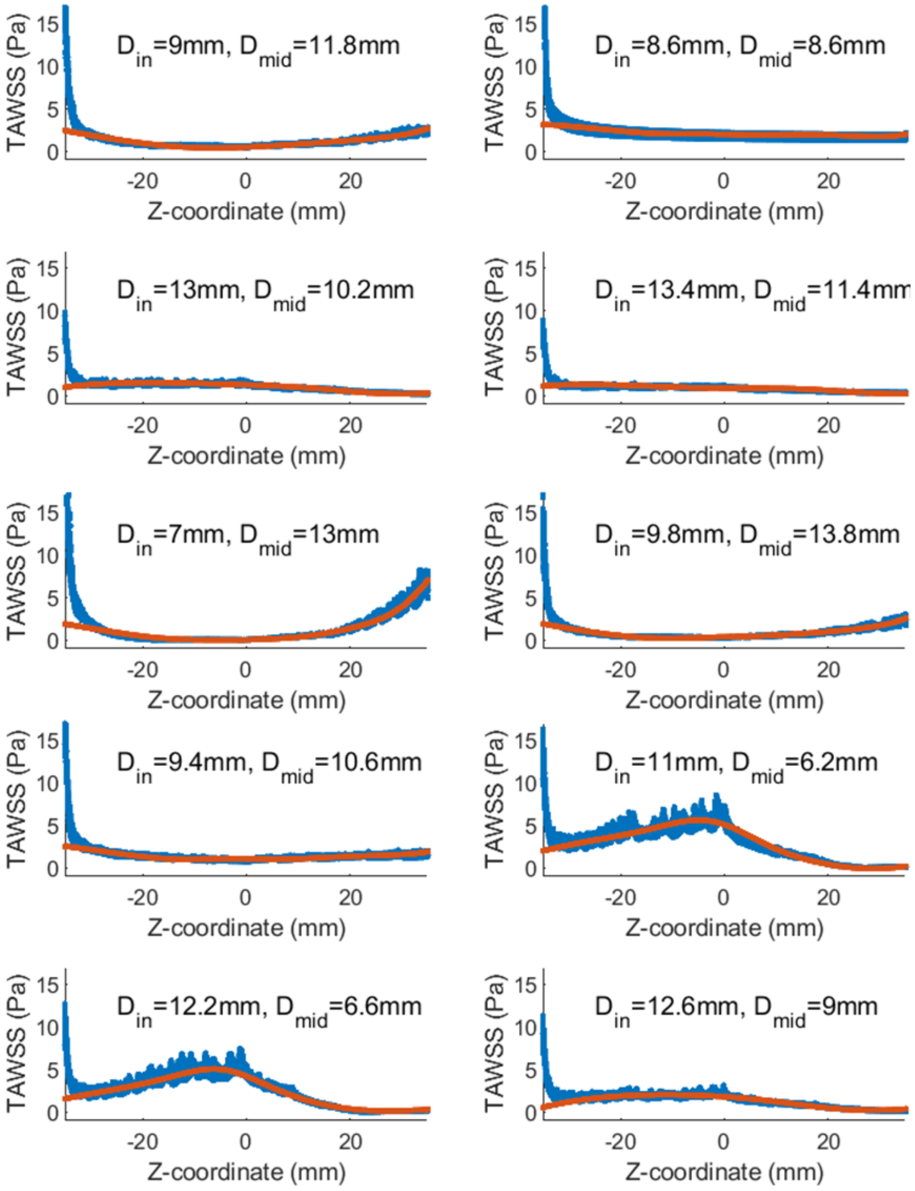


Figure 5: Results of the surrogate model training for the carotid validation case. The TAWSS from the CFD simulation (blue) and the ANN (red) is shown in function of the Z-coordinate for the ten first combinations of $D_{in}$ and $D_{mid}$.

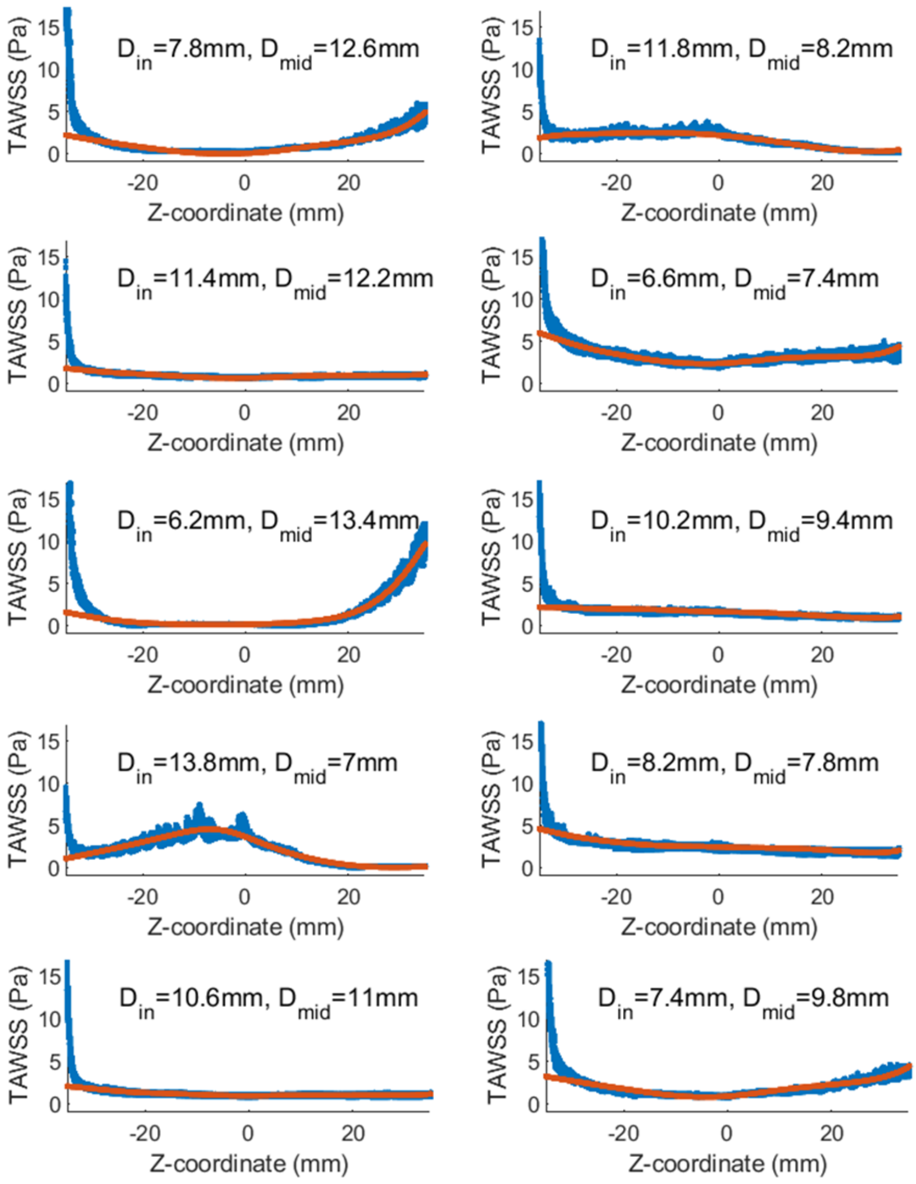


Figure 6: Results of the surrogate model training for the carotid validation case. The TAWSS from the CFD simulation (blue) and the ANN (red) is shown in function of the Z-coordinate for the ten last combinations of $D_{in}$ and $D_{mid}$.

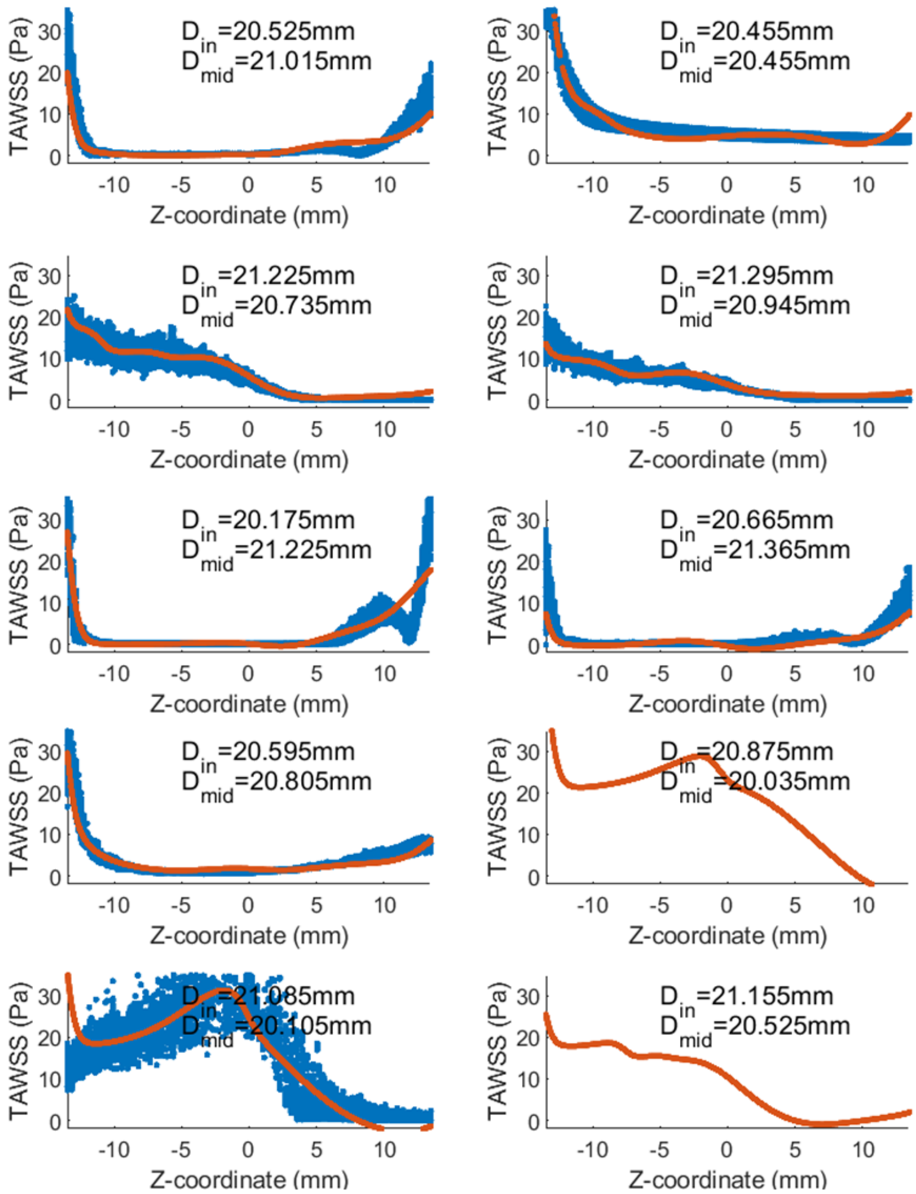


Figure 7: Results of the surrogate model training for the pulmonary validation case. The TAWSS from the CFD simulation (blue) and the ANN (red) is shown in function of the Z-coordinate for the ten first combinations of $D_{in}$ and $D_{mid}$. No stable CFD solution could be found for the case with $D_{in}$=20.875mm and $D_{mid}$=20.035mm, and the case with $D_{in}$=21.155mm and $D_{mid}$=20.525mm. These cases were hence not taken into account for the training of the ANN.

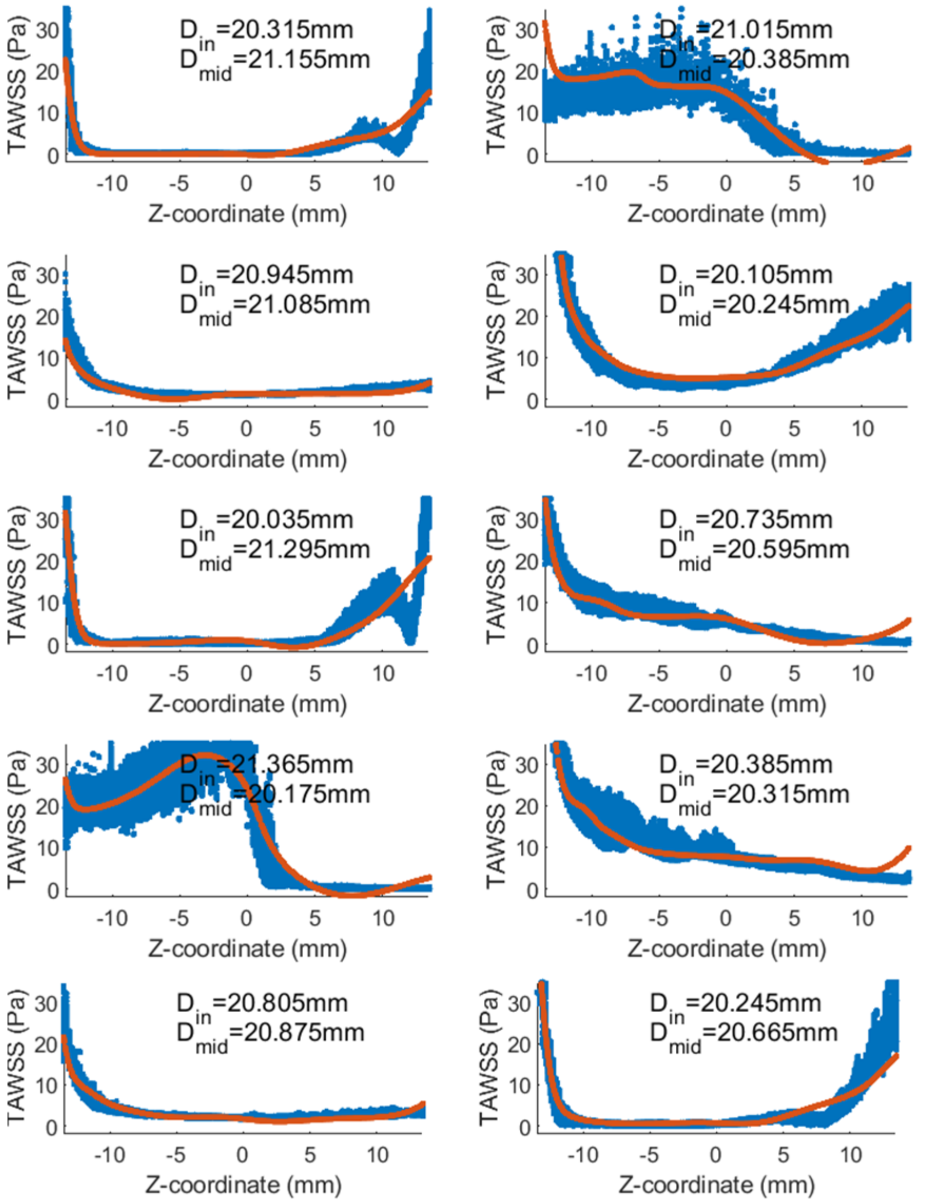


Figure 8: Results of the surrogate model training for the pulmonary validation case. The TAWSS from the CFD simulation (blue) and the ANN (red) is shown in function of the Z-coordinate for the ten last combinations of $D_{in}$ and $D_{mid}$.
